# PGC1α Regulates the Endothelial Response to Fluid Shear Stress *via* Telomerase Reverse Transcriptase Control of Heme Oxygenase-1

**DOI:** 10.1101/2021.05.26.445830

**Authors:** Shashi Kant, Khanh-Van Tran, Miroslava Kvandova, Amada D. Caliz, Hyung-Jin Yoo, Heather Learnard, Siobhan M. Craige, Joshua D. Hall, Juan M. Jiménez, Cynthia St. Hilaire, Eberhard Schulz, Swenja Kröller-Schön, John F. Keaney

## Abstract

Fluid shear stress (FSS) is known to mediate multiple phenotypic changes in the endothelium. Laminar FSS (undisturbed flow) is known to promote endothelial alignment to flow that is key to stabilizing the endothelium and rendering it resistant to atherosclerosis and thrombosis. The molecular pathways responsible for endothelial responses to FSS are only partially understood. Here we have identified peroxisome proliferator gamma coactivator-1α (PGC-1α) as a flow-responsive gene required for endothelial flow alignment *in vitro* and *in vivo*. Compared to oscillatory FSS (disturbed flow) or static conditions, laminar FSS (undisturbed flow) increased PGC-1α expression and its transcriptional co-activation. PGC-1α was required for laminar FSS-induced expression of telomerase reverse transcriptase (TERT) *in vitro* and *in vivo* via its association with ERRα and KLF4 on the *TERT* promoter. We found that TERT inhibition attenuated endothelial flow alignment, elongation, and nuclear polarization in response to laminar FSS *in vitro* and *in vivo*. Among the flow-responsive genes sensitive to TERT status was heme oxygenase-1 (HMOX1), a gene required for endothelial alignment to laminar FSS. Thus, these data suggest an important role for a PGC-1α-TERT-HMOX1 axis in the endothelial stabilization response to laminar FSS.

## Introduction

The endothelium exerts considerable control over vascular homeostasis with important roles governing vascular tone, inflammation, and metabolism ^1,2^. Normal endothelial function is characterized by a quiescent cell phenotype that is non-proliferative, non-migratory, and exhibits a cell surface that prevents thrombosis, inflammation, and lipid deposition, thereby resisting atherosclerosis and vascular disease ^3,4^. A key stabilizing stimulus for endothelial quiescence is laminar fluid shear stress (also called undisturbed flow) passing over the cell surface, a common feature of straight vascular segments with little to no curvature ^5^. Endothelial cells reorient and change their shape in order to align their long axis to physiological fluid shear stress (FSS) ^6^. In contrast, curved and branched arteries experience multidirectional and chaotic FSS (also called disturbed flow). Areas with disturbed flow promote a less stable, activated, and dysfunctional endothelial phenotype that is more susceptible to inflammation and atherosclerosis ^5,7-11^. Thus, endothelial response to fluid shear stress is a critical element of vascular homeostasis.

Endothelial cell responses to fluid shear stress involve both mechanotransduction of the flow signal and coordinated regulation of signaling pathways and gene expression that dictate the phenotypic consequences. A number of mechanosensors and mechanotransducers have been implicated in flow sensing, including ion channels, PECAM-1, G protein-coupled receptors, junctional proteins, VEGF receptors, and even primary cilia ^5,7,12,13^. A number of pathways have been implicated in the phenotypic response to physiologic vs. pathologic flow. When exposed to low flow rate or disturbed flow NF-κB, Yap/Taz, β-catenin and Smad2/3 have high activity, where they cooperate to promote remodeling, which includes activation of inflammatory mediators and, to variable extents, induction of endothelial to mesenchymal transition ^14-16^. Physiological laminar flow appears to activate KLF2/4, Notch, and Alk1-Smad1/5 that induce genes known to contribute to vascular stability ^17-19^. In particular, KLF2 and KLF4 have been implicated in promoting vasodilation and the inhibition of both inflammation and thrombosis ^20^. Among the genes regulated by KLF2 and KLF4 is NOS3 that codes for the endothelial isoform of nitric oxide synthase (eNOS) ^20^. This enzyme contributes importantly to the vascular homeostatic environment by promoting vasodilation and limiting atherothrombosis. We recently reported that the transcriptional coactivator, peroxisome proliferator gamma coactivator-1a (PGC1α), was an important determinant of eNOS expression *in vitro* and *in vivo* ^21^. Here we sought to determine if PGC1α and its downstream targets play a role in endothelial response to fluid shear stress.

## Result

### PGC1α is needed for endothelial cell response to fluid shear stress

Endothelial cells change their shape and reorient themselves by aligning in the direction of flow accompanied by nuclear polarization and elongation in response to physiological levels of fluid shear stress ^6^. To examine the impact of flow on endothelial alignment, we examined the response of HAECs to oscillatory vs. laminar FSS (Disturbed flow vs. undisturbed flow). As expected, oscillatory FSS yielded random orientation of HAECs in culture, whereas laminar FSS produced HAECs alignment in the direction of flow (Supplemental Figure. 1A and 1B). Laminar FSS also led to increased expression of key flow-responsive genes, including *KLF2* ^18^, *KLF4* ^19,22^, *HMOX1* ^23^, and *HEY1* ^24^ (Supplemental Figure. 1C-D). Since we and others have found that PGC1α has implications for endothelial function ^21,25^, we examined its expression as a function of FSS. We found that PGC1α protein and mRNA levels were significantly higher after atheroprotective laminar shear stress (LSS) compared to static control (Figure. 1A-B). Similarly, PGC1α mRNA was upregulated with laminar compared to oscillatory flow in two different human endothelium cell types (Figure. 1C). These data suggest that PGC1α may play an important role during physiological fluid shear stress.

**Figure 1.**
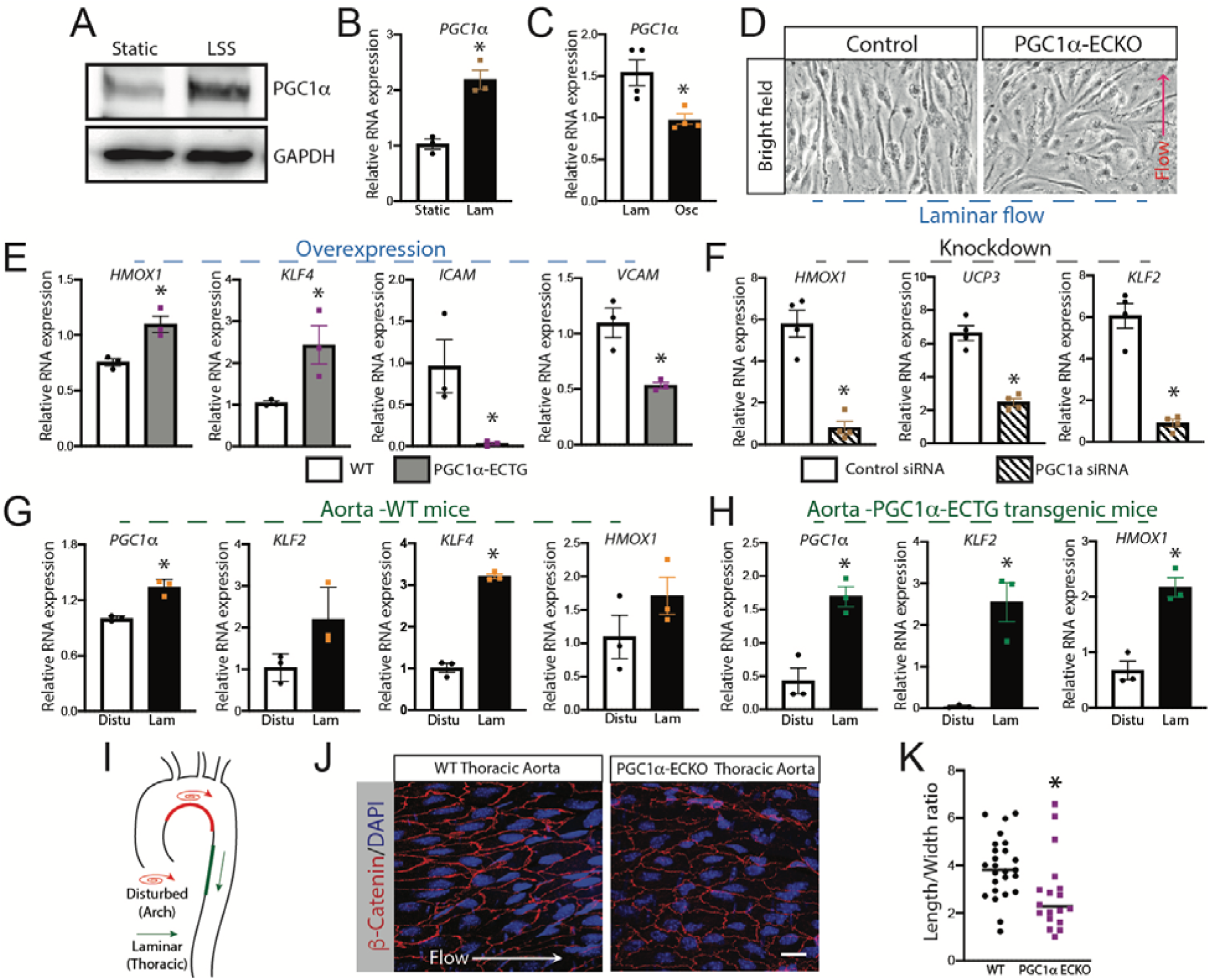
PGC1a regulates endothelial cell function during flow shear stress. A-B) PGC1α protein expression (A) or mRNA expression (B) from HUVECs were measured by either western blots or RT-qPCR after cells were subjected to either static or laminar flow shear stress for 48 hours. C) PGC1α mRNA expression measured by RT-qPCR from HUVECs after exposure of 48 hours of laminar or oscillatory FSS. D) Bright-field image of Mouse lung endothelial cells (MLECs) isolated from either wild-type (WT) or PGC1α ECKO mice after exposure to laminar FSS. E) MLECs were isolated from WT and PGC1α-ECTG mice, and RT-qPCR were performed for different genes related to endothelial function after exposure to laminar FSS. F) HAECs were either treated with scrambled or *PGC1α* siRNA, and RT-qPCR was performed for different genes related to endothelial function after exposure to laminar FSS for 48 hours. G-H) Aortae were isolated from either WT (G) or PGC1α-ECTG (H) mice and mRNA was isolated, and RT-qPCR was done for the arch and thoracic region for disturbed and laminar flow. (I) Sample sites of disturbed (oscillatory) vs. laminar fluid shear stress in mouse aorta. (J) *En face* staining with β-Catenin and DAPI in WT and PGC1α -ECKO aorta. Scale bar, 20Lμm. (K) Composite data of length/width ratio of the endothelium in the thoracic region of mouse aortae. *n*□=□3 – 6 in each group. All the experiments were repeated 3 – 5 times. Statistically significant differences between groups are indicated (\**P*□<□0.05 by Student’s *t-test*). The data are mean□±□SEM. Scale bar, 5□μm.

To probe the role of PGC1α directly, we utilized the molecular strategy of loss of function and gain of function by using PGC1α knockout and overexpression mouse models of PGC1α. We isolated mouse lung endothelial cells (MLECs) from WT or PGC1α endothelial cell specific knockout mice (PGC1α-ECKO) mice. Wild-type MLECs subjected to laminar FSS for 48 hours exhibited flow alignment, whereas PGC1α ECKO MLECs did not (Figure. 1D). Next, we used a gain of function strategy with MLECs from our endothelial specific PGC1α overexpressing transgenic mice (PGC1α-ECTG) that exhibit ∼60% increase in endothelial PGC1α compared to WT MLECs ^21^. We found that that static PGC1α-ECTG MLECs exhibited upregulation of the flow-responsive genes *Klf4* ^8,19^ and *Hmox1* ^23^ compared to wild-type cells (Figure. 1E).

Moreover, MLEC PGC1α gain-of-function suppressed *Icam-1* and *Vcam-1* mRNA expression (Figure. 1E). In HAECs treated with PGC1α siRNA, we observed blunted upregulation of *HMOX1* and *KLF2* ^18^ in response to laminar FSS compared to the scrambled siRNA control (Figure. 1F).

To determine the physiologic role of PGC1α, we examined intact aortic segments from areas of disturbed (inner arch) and laminar (descending thoracic) flow as in Figure. 1I. We observed relatively greater PGC1α mRNA expression in the laminar flow segments compared to areas of disturbed flow (Figure. 1G-H). Similarly, mRNA expression of *Hmox1, Klf2*, and *Klf4* were similarly upregulated in the laminar vs. disturbed flow regions of the aorta in WT and PGC1α-ECTG mice (Figure. 1G-H). Next, we used *en face* staining of the aorta with the endothelial junction marker β-catenin, to assess morphological endothelial responses to fluid shear stress. Endothelial cells adapt to laminar FSS by exhibiting planar cell polarity with the flow ^26,27^. The endothelial length to width ratio is a key index of flow alignment and planar cell polarity ^28^. Qualitative assessment of *en* face staining demonstrated a highly elongated polygons endothelial cell shape and orientation to the vessel’s axis in descending aorta (Figure. 1J-K). Objectively, we observed a reduced endothelial length to width ratio in PGC1α-ECKO descending aorta compare to WT littermate controls (Figure. 1J-K). Collectively these data suggest that PGC1α is a crucial element of the endothelial cell response to fluid shear stress in terms of both morphology and the expression of FSS-responsive genes.

### Endothelial PGC1α is required for the endothelial response to exercise

Prolonged exercise is known to increase vascular FSS to the high physiological range and is associated with upregulation of key FSS-responsive genes that include *NOS3, KLF2/4, and HMOX1* ^29-31^. In WT mice subjected to chronic exercise, PGC1α mRNA expression in the aorta was significantly greater than sedentary animals (Figure. 2A). We also observed that *Hmox1*, as well as telomerase reverse transcriptase (*Tert)*, a gene sensitive to PGC1α status in vascular smooth muscle ^32^, were upregulated in WT mice with chronic exercise mice, but not in PGC1α-ECKO mice (Figure. 2A). Since PGC1α is known to contribute to reactive oxygen species (ROS) detoxification ^33^, we examined ambient vascular ROS as a function of exercise. Superoxide, mitochondrial ROS, and H_2_O_2_ were estimated by dihydroethidium (DHE), MitoSOX, and Amplex Red, respectively. We found that chronic exercise reduced all three indices of ROS; however, this effect was lost in PGC1α -ECKO animals (Figure. 2B). Endothelial function determined as aortic relaxation to acetylcholine (Ach) improved with chronic exercise (Figure. 2C), but this effect was lost in PGC1α -ECKO animals. These data implied that PGC1α is an important component of endothelial function during exercise and is required for full exercise-induced *Hmox1* and *Tert* upregulation.

**Figure 2.**
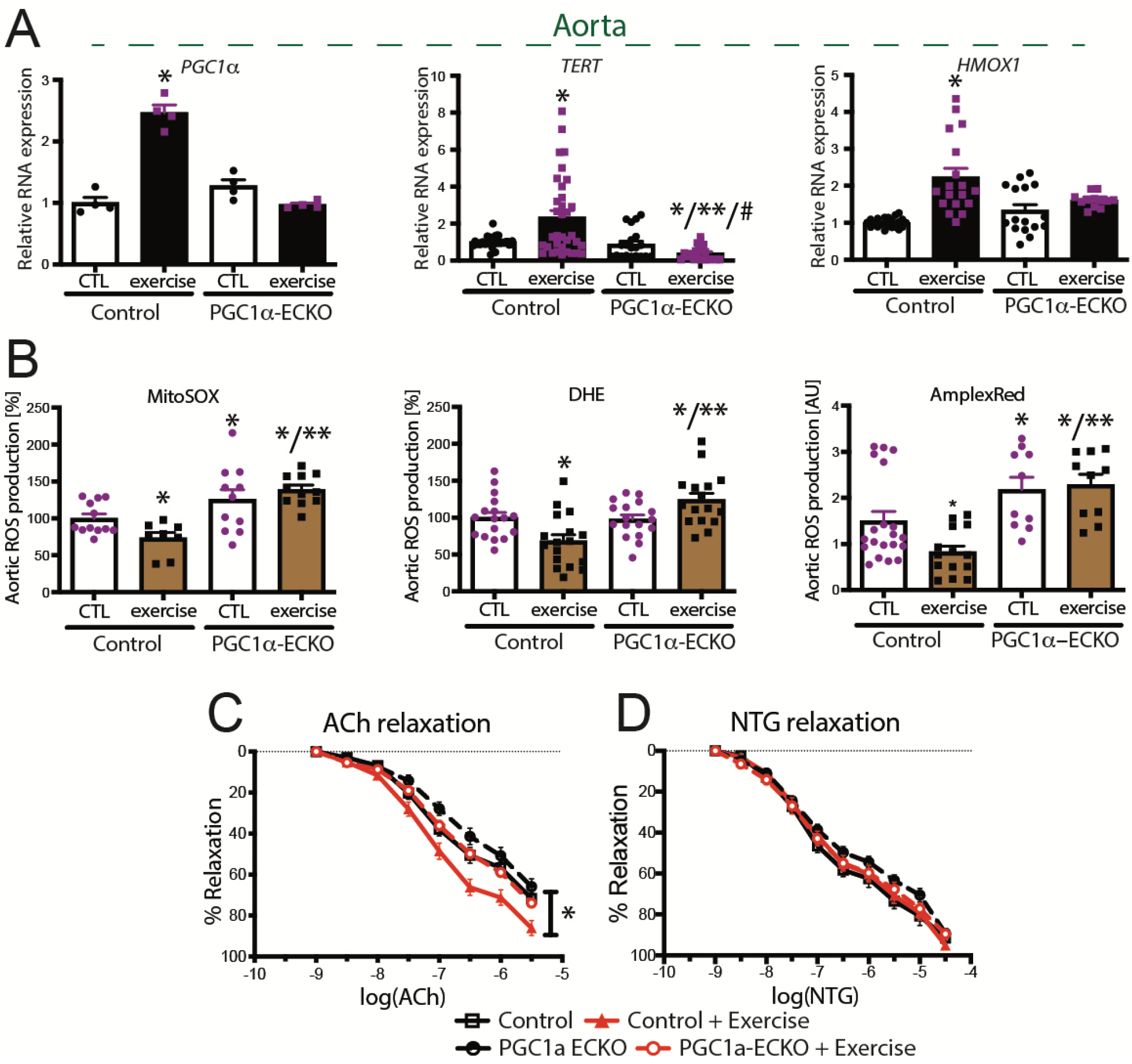
PGC1α is required for endothelial function during exercise. A) Aortae were isolated from either WT or PGC1α ECKO mice and mRNA was measured by RT-qPCR for indicated genes before and after exercise. B) Reactive oxygen species (ROS) production was measured as indicated in WT and PGC1α ECKO mice before and after exercise. C-D) Endothelial function was measured by aortic isometric force in response to acetylcholine (Ach; C) and nitroglycerin (NTG; D) in organ chamber assay. N□=□4–30/group, \**P*□<□0.05 vs. control mice.

### PGC1α impacts FSS-dependent HMOX1 expression via ERRα and KLF4

PGC1α impacts gene expression by the co-activation of transcription factors ^34^. The transcription factors, ERRα and KLF4, play a critical role in regulating endothelial function ^21,35^. In cardiac myocytes, ERRα and KLF4 interact with each other and can form a complex with the PGC1α protein ^36^. Therefore, we examined the role of ERRα and KLF4 in PGC1α– mediated endothelial responses to laminar FSS. First, using siRNA directed against ERRα in HAECs, we observed that ERRα controls endothelial upregulation of *HMOX1* expression in response to laminar FSS (Figure. 3A). Similarly, *KLF4* siRNA inhibited *HMOX1* expression in the setting of laminar FSS (Figure. 3B). In contrast, KLF4 and ERRα had no reciprocal effect on each other’s expression with laminar FSS (Figure. 3A-B). As *HMOX1* expression depends on both ERRα and KLF4, we tested their interaction in endothelial cells. ERRα immunoprecipitation demonstrated that it forms and exists in a complex with PGC1α and KLF4 in human endothelial cells (Figure. 3C). These data suggest that ERRα with KLF4 plays an important role in PGC1α regulated HMOX1 regulation.

**Figure 3.**
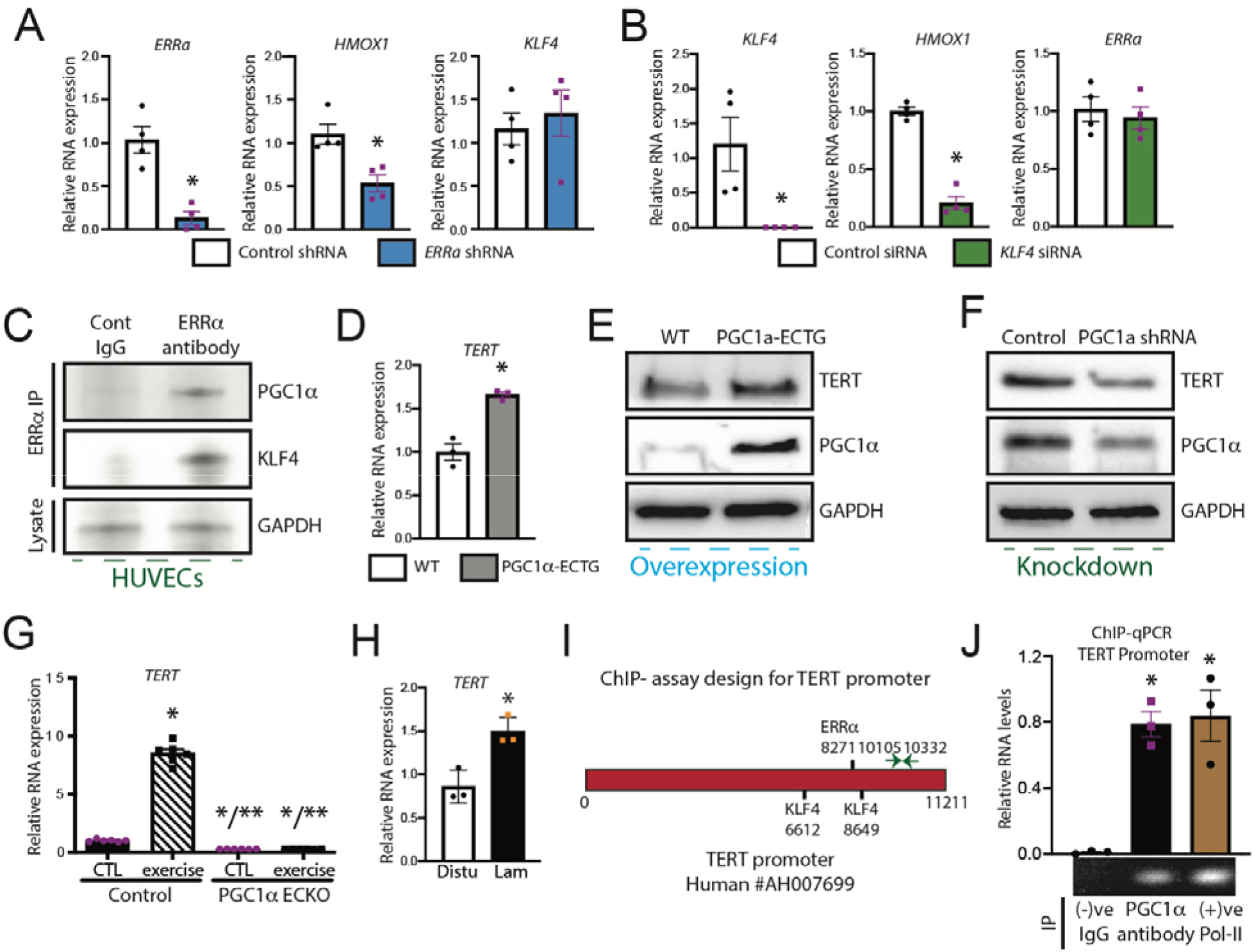
PGC1α regulates TERT expression. A-B) HAECs were either treated with scrambled or *ERRα* siRNA (A) or scrambled or *KLF4* siRNA (B), and RT-qPCR was performed for different shear stress-related genes after exposure of cells with laminar FSS for 48 hours. C) HUVECs were lysed and immunoprecipitation (IP) was performed either with control IgG or ERRα antibody and immunoblotting was done with antibodies against PGC1α and KLF4. Lysates were examined by probing with GAPDH antibody. D-E) MLECs were isolated from control and PGC1α -ECTG mice, and either RT-qPCR (D) or immunoblot analysis (E) were performed with the probes and antibodies as indicated. F) Lysates prepared from HAECs treated with control shRNA or shRNA against *PGC1α* (48□hrs) were examined by immunoblot analysis using antibodies for TERT, PGC1α and GAPDH. G) MLECs were isolated from WT and PGC1α ECKO mice aortae and mRNA expression was measured by RT-qPCR for *TERT* gene before and after exercise. H) Aortae were isolated, and mRNA expression was measured for TERT by RT-qPCR in the arch and thoracic region of WT mice. I-J) ChIP-qPCR analysis of PGC1α recruitment to the TERT promoter region was performed in HUVECs. *N*□=□3 - 4 in each group. All the experiments were repeated 3 – 6 times. Statistically significant differences between groups are indicated (\**P*□<□0.05 by Student’s *t-test*). The data are mean□±□SEM.

### PGC1α dictates telomerase reverse transcriptase expression in the endothelium

One identified PGC1α-dependent gene is telomerase reverse transcriptase (*TERT*) that, along with the telomerase RNA component (*TERC*), forms the telomerase complex that has been implicated in vascular aging ^32^. In cultured endothelial cells, oxidative stress stimulates nuclear export of TERT to mitochondria as a protective mechanism ^37^, and inhibition of telomerase impairs flow-mediated nitric oxide bioactivity in human arterioles ^38^ However, the role of TERT in endothelial responses to FSS is incompletely understood. Therefore, we used PGC1α gain- and loss-of-function to probe its implications for TERT in FSS responses. We found that PGC1α - ECTG MLECs exhibited upregulation of Tert mRNA and protein (Figure. 4D-E), whereas PGC1α shRNA impaired endothelial TERT protein expression (Figure. 3F). Next, we probed TERT expression as a function of FSS. Similar to the PGC1α exercise-induced FSS upregulated aortic Tert mRNA in WT mice, but not in PGC1α-ECKO mice (Figure. 3G). Furthermore, we found that similar to PGC1α, TERT expression is higher in the laminar flow (thoracic) region of the aorta than in the disturbed flow (inner arch) region (Figure. 3H). Next, to find how PGC1α controls the expression of TERT mRNA, we did a chromatin immunoprecipitation (ChIP) assay for the TERT promoter with PGC1α antibody and found that PGC1α binds to the TERT promoter region (Figure. 3I-J). These data clearly showed that PGC1α is required for TERT expression.

**Figure 4.**
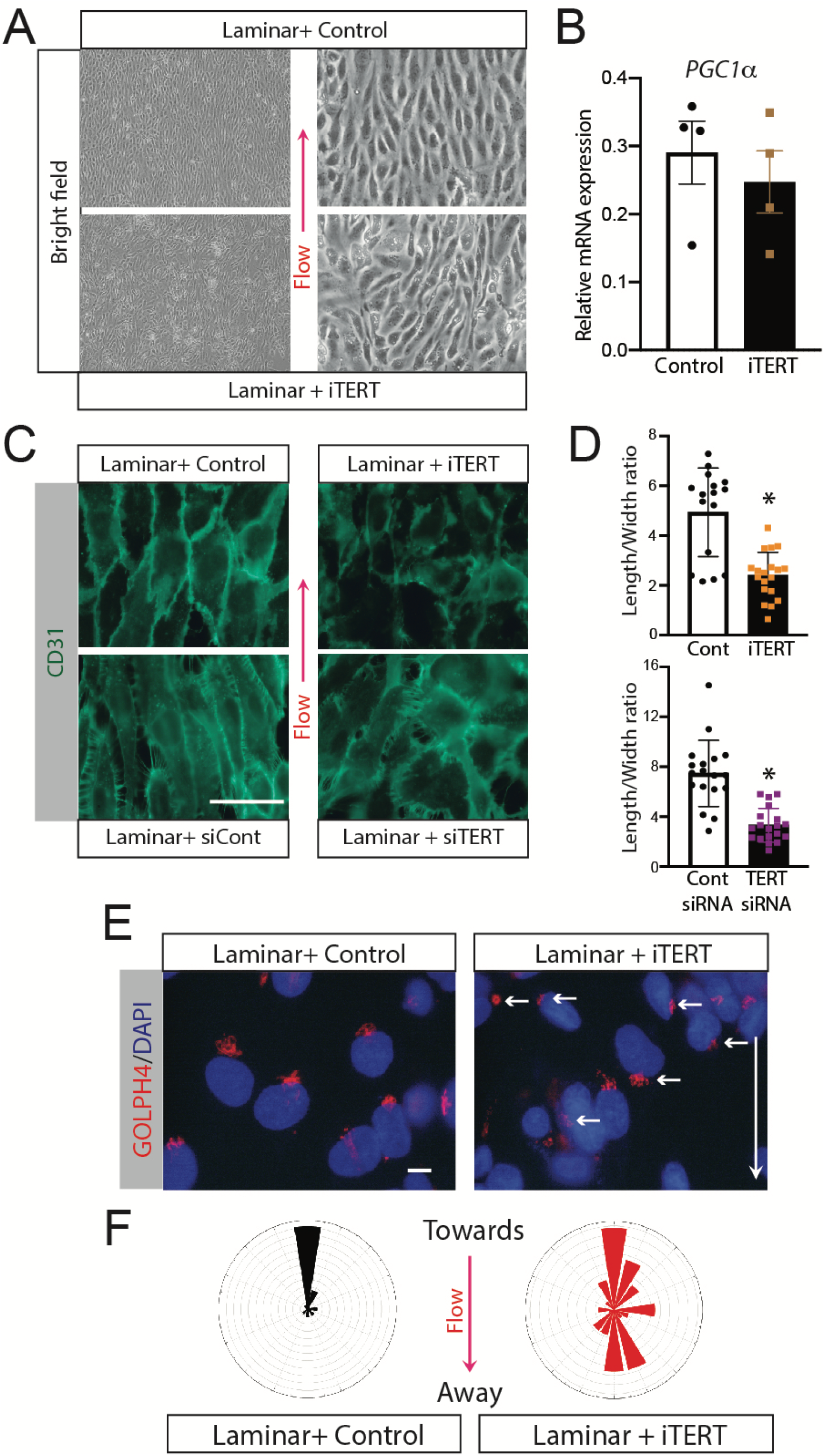
TERT is required for endothelial alignment to the flow. A) Bright-field image of HAECs after exposure of laminar FSS in the presence of either control or TERT inhibitor. B) mRNA was isolated, and expression was measured for *PGC1α* expression by RT-qPCR treated by either control or TERT inhibitor. C) HAECs morphology was measured by staining for CD31/DAPI as a function of pharmacologic (iTERT) or genetic (*siTERT*) inhibition of TERT. Scale bar, 25Lμm. D) Length to width ratio of CD31 stain HAECs measured after TERT inhibition in the presence of laminar FSS. E) HAECs, length/width ratio was measured after staining of CD31 with or without TERT inhibitor. E) HAEC nuclear polarization to Golgi towards laminar flow direction was measured with GOLPH4 (Golgi) and DAPI (nuclei) staining with and without TERT inhibitor. Scale bar, 10Lμm. F) Compass plots of Golgi/nuclear angle as a function of TERT inhibition. Each ring represents an observation of an average of different fields of control or TERT inhibitor-treated cells. All the experiments have been repeated 3 – 6 times. Statistically significant differences between groups are indicated (\**P*L<L0.05 by Student’s *t-test*). The data are meanL±LSEM. Scale bar, as indicated.

### TERT is required for endothelial alignment, elongation, and polarization

We examined the impact of TERT on endothelial cell responses to FSS using both pharmacologic and molecular approaches. Endothelial cells treated with the TERT inhibitor, BIBR1532 (iTERT) ^39^, exhibited impaired orientation and alignment in the direction of laminar FSS compared to vehicle-treated cells (Figure. 4A). Pharmacologic TERT inhibition had no impact on PGC1α expression (Figure. 4B). Next, we treated HAECs with siRNA directed against *TERT* (siTERT) and observed ∼80% reduction in *TERT* expression levels (Supplemental Figure. 2). Suppression of *TERT* expression also inhibited HAECs visual alignment to laminar FSS compared to scrambled siRNA control (Figure. 4C). This lack of alignment was also manifest as reduced length: width ratio in response to laminar FSS in the presence of either pharmacologic or molecular TERT inhibition (Figure. 4D). Another endothelial response to laminar FSS is nuclear polarization with the Golgi directed against the flow ^40^. Therefore, we examined this phenomenon using the Golgi marker, GOLPH4. We found that TERT inhibition prevented nuclear polarization towards laminar flow (Figure. 4E-F). Collectively, these data show that TERT plays an important role in endothelial cell alignment and polarization in response to laminarFSS.

### TERT is required for mitochondrial responses to FSS

PGC1α plays an important role in mitochondrial function and biogenesis ^41-43^. TERT is known to have non-canonical functions that include translocation to the mitochondria and mitochondrial stabilization with oxidative stress ^37,44^. We probed the implications of TERT in the endothelial FSS response by exposing HAECs to oscillatory vs. laminar FSS in the presence or absence of TERT inhibition. We found that laminar FSS produced elongated and branched mitochondrial staining with MitoTracker (Figure. 5A). In contrast, TERT inhibition produced punctate mitochondrial morphology that was reminiscent of oscillatory flow (Figure. 5A). Composite data also indicate that TERT inhibition with laminar FSS produces an endothelial response that mimics oscillatory flow with respect to mitochondrial mass by MitoTracker staining and network formation by mitochondrial length (Figure. 5B). To determine the role of TERT on mitochondrial ROS, we stained laminar FSS exposed HAECs with MitoSOX as a function of TERT inhibition. Compared to vehicle-treated cells, TERT inhibition enhanced the mitochondrial ROS signal (Figure. 5C). TERT has been implicated in telomere length maintenance. Although these experiments are short-term, we did examine telomere length over the time course of our experiment and found that none of the conditions used for TERT inhibition in our experiments had any impact on telomere length (Supplemetal Figure. 3). These data imply that TERT plays an important role in the endothelial mitochondrial response to laminar FSS that is independent of its canonical function on telomere length. To determine if these findings play a role *in vivo*, we used Tert knockout mice. *En face* staining of endothelial borders with β-catenin in laminar flow segments as in Figure. 1 revealed qualitatively impaired flow alignment in Tert knockout segments vs. wild-type (Figure. 5D). Composite data on the length:width ratio revealed significantly reduced elongation (Figure. 5E). These data strongly suggest that TERT is required for optimal endothelial response to laminar FSS *in vivo*.

**Figure 5.**
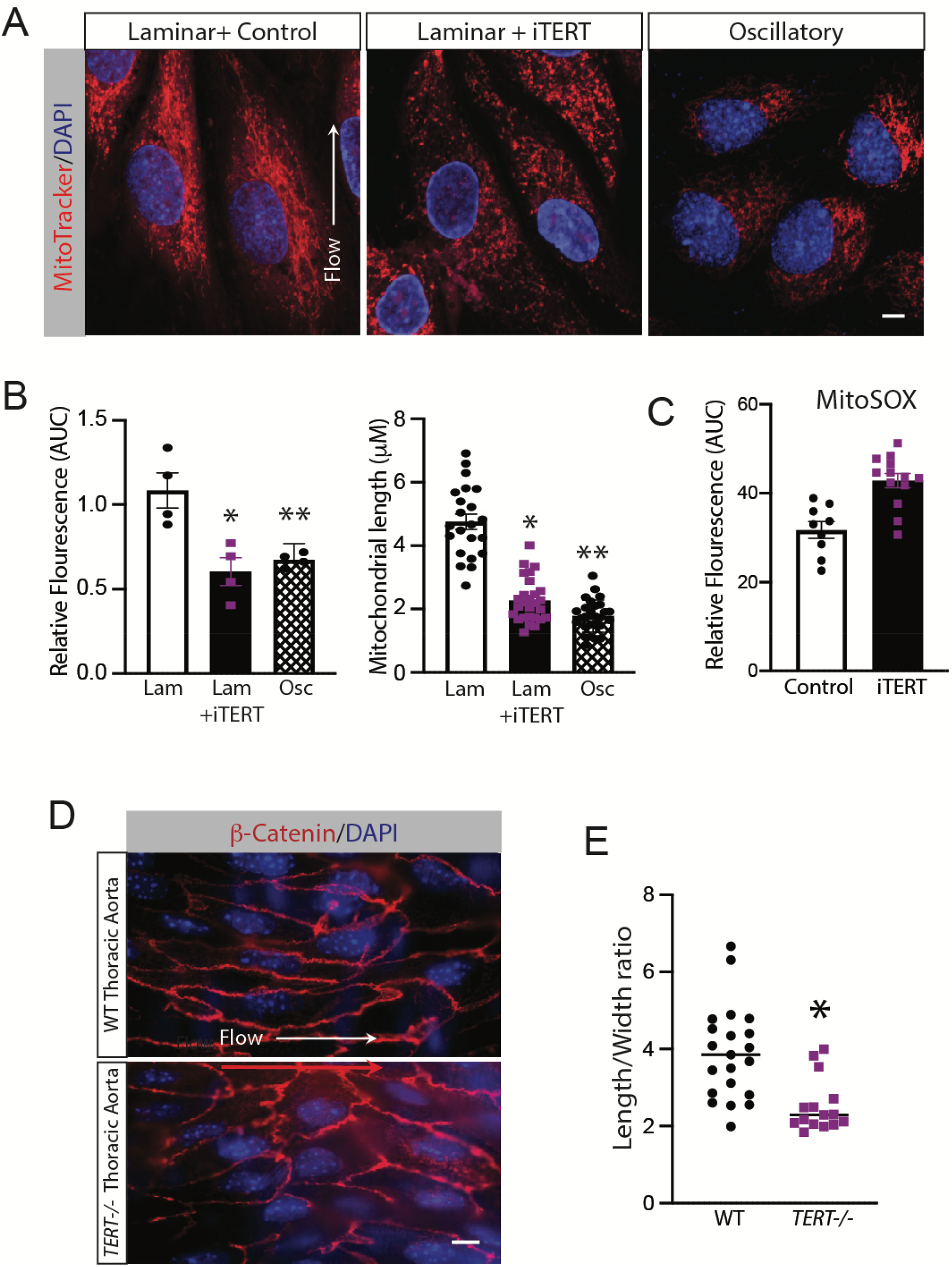
TERT regulates mitochondrial structure and function. A) Mitochondrial morphology and mass (MitoTracker fluorescence) were measured in the presence of Laminar flow with and without TERT inhibition (iTERT-BIBR1532) and compared with the oscillatory flow. Scale bar, 5Lμm. B) Quantification of mitochondrial mass and morphology during FSS. C) ROS production in HAECs either treated with control or TERT inhibitor. D) *En face* staining with β-Catenin and DAPI in WT and TERT knockout aortae. Scale bar, 10Lμm. E) Composite data of length/width ratio of the endothelium in WT and TERT knockout aortae. *n*L=Las indicated in each group. All the experiments have been repeated 3 – 6 times. Statistically significant differences between groups are indicated (\**P*L<L0.05 by Student’s *t-test*). The data are meanL±LSEM. Scale bar, as indicated.

### TERT is required for normal laminar FSS-induced HMOX1 expression

Since our data indicate that both PGC1α (Figure. 1F) and ERRα (Figure. 3A) participate in laminar FSS-induced HMOX1 expression, we tested the role of TERT in this process. As expected, both *HMOX1* and *KLF4* were upregulated with laminar FSS (Figure. 6A). Moreover, endothelial overexpression of PGC1α increased HMOX1 protein under static conditions (Figure. 6B), whereas siRNA against ERRα attenuated HMOX1 protein during laminar flow (Figure. 6C). Consistent with these findings, exercise-induced FSS upregulated aortic *Hmox1* expression in a manner dependent upon endothelial PGC1α (Figure. 6D). In terms of Tert, we observed that compared to oscillatory FSS, laminar FSS upregulated Hmox1 protein in a manner dependent upon TERT activity as it was inhibited by a TERT specific inhibitor BIBR1532 (Figure. 6E). *HMOX1* expression appears to be downstream of *PGC1α/TERT/KLF2* as *HMOX1*-directed shRNA had no impact on laminar FSS-mediated expression of these genes (Figure. 6F).

**Figure 6.**
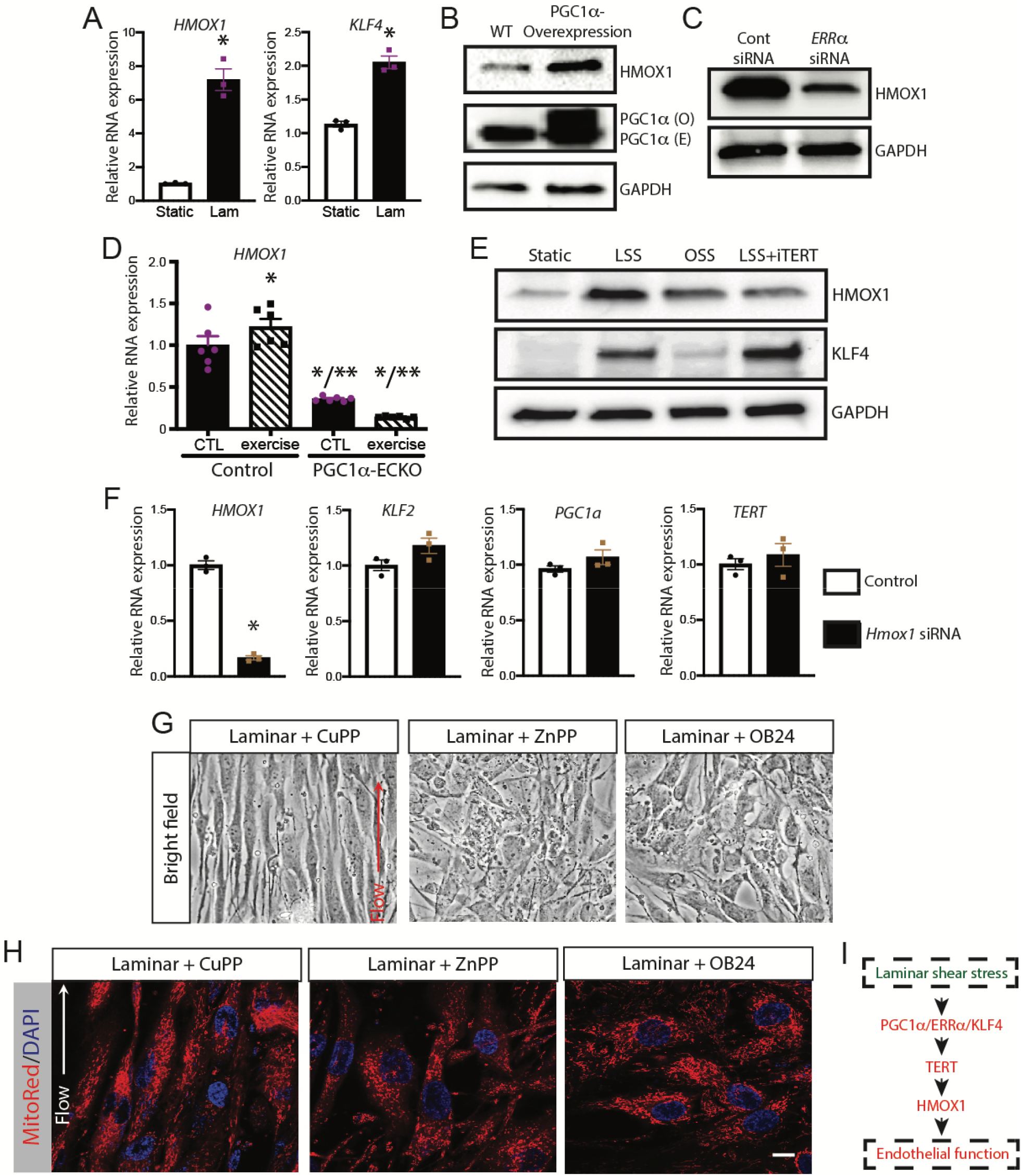
PGC1a-TERT regulates HMOX1 expression. A) *HMOX1* and *KLF4* mRNA expression in HAECs after LSS. B) MLECs were isolated from control and PGC1α-ECTG mice and immunoblot analysis was performed with the antibodies as indicated. C) Lysates prepared from HAECs after treatment with either control or *ERR siRNA* (48□hrs) were examined by immunoblot analysis using HMOX1 and GAPDH antibodies. D) MLECs were isolated from WT, and PGC1α ECKO mice aortae and mRNA expression was measured by RT-qPCR for the *HMOX1* gene before and after exercise. E) HUVECs were exposed with laminar or oscillatory FSS with or without TERT inhibitor and immunoblot analysis was performed with the antibodies as indicated. F) HAECs were either treated with scrambled or *HMOX1* siRNA and RT-qPCR was performed for different genes related to endothelial function after the exposure of laminar FSS for 48 hours. G) Bright-field image of HAECs after exposure of laminar FSS in the presence of either control (CuPP 1mM) or two different HMOX1 inhibitors (ZnPP 1mM and OB 24 hydrochloride 1mM). H) Cell and mitochondrial morphology (MitoRed fluorescence) were imaged in the presence of Laminar flow with and without HMOX1 inhibition. I) D) Schematic diagram of LSS-induced PGC1α-TERT-HMOX1 pathway. Scale bar, 5□μm. *n*□=□3 - 6 in each group. All the experiments were repeated 3 – 5 times. Statistically significant differences between groups are indicated (\**P*□<□0.05 by Student’s *t-test*). The data are mean□±□SEM.

### HMOX1 is required for the endothelial cell response to the laminar flow

To examine if HMOX1 dictates endothelial FSS responses, we used a pharmacological approach by using two different inhibitors zinc (II) protoporphyrin IX (ZnPP) and 1-[[2-[2-(4-Bromophenyl)ethyl]-1,3-dioxolan-2-yl]methyl]-1*H*-imidazole hydrochloride (OB 24 hydrochloride) for HMOX1 ^45-47^. We used copper protoporphyrin IX (CuPP) as control. CuPP is a similar compound to ZnPP but does not inhibits HMOX1 ^45^. Both HMOX1 inhibitors, ZnPP and OB 24 hydrochloride, impaired HAECs flow alignment along the direction of laminar FSS compared to control, copper protoporphyrin IX (CuPP) treated cells (Figure. 6G). Similarly, HMOX1 inhibition also impaired mitochondrial network formation in response to laminar FSS, leaving HAECs with punctate mitochondria as determine by MitoRed staining (Figure. 6H).

Exposure to low flow rate or disturbed flow induction of endothelial to mesenchymal transition leads to endothelial dysfunction^14-16^. To determine the role of the PGC1α -TERT-HMOX1 pathway from endothelial to mesenchymal transition, we used RT-qPCR to check the expression of mesenchymal marker into MLECs isolated from PGC1α-ECTG mice as well as HAECs cells treated with TERT and HMOX1 siRNA. We found that forced overexpression of PGC1α in MLECs suppresses the mesenchymal markers SM22α (*Tagln*), Sca1 (*Atxn1*), MMP2 and ICAM1 (Supplemetal Figure. 4A). Furthermore, similar to PGC1α result mesenchymal markers were upregulated in HAECs with either TERT siRNA or HMOX1 siRNA compare to control (Supplemetal Figure. 4B-C). These data support a cascade depicted in Figure. 6I whereby PGC1α/ERRα contributes to laminar FSS-induced TERT expression that is needed for the HMOX1 upregulation required for full endothelial FSS responsiveness.

## Discussion

The data presented here indicate that PGC1α is a key element of the endothelial response to fluid shear stress. In the absence of PGC1α, endothelial cells exhibited an impaired ability to align and elongate as well as the upregulation of key genes characteristic of the response to laminar FSS. These findings appear relevant *in vivo* as aortic segments with laminar FSS exhibited PGC1α upregulation, and animals lacking endothelial PGC1α demonstrated impaired flow alignment in laminar flow sections of the aorta. Similarly, exercise-induced FSS was associated with improved aortic endothelial function, antioxidant defenses, and reduced ROS that were all impaired in the absence of endothelial PGC1α. We found that PGC1α exerted its influence via a complex, including ERRα and KLF4, that afford PGC1α binding to the TERT promoter to increase TERT expression. In keeping with these findings, TERT upregulation was required for the action of PGC1α as TERT loss-of-function recapitulated the endothelial PGC1α loss-of-function phenotype, including endothelial elongation, flow alignment, mitochondrial stabilization, and Hmox1 upregulation with laminar FSS. With regard to the latter, we found that HMOX1 loss-of-function was a key downstream mediator of endothelial PGC1α as HMOX1 inhibition reproduced the endothelial PGC1α-null phenotype, including flow alignment, elongation, and mitochondrial stabilization. Taken together, these data identify a PGC1α-TERT-HMOX1 axis as a key element of endothelial cell responses to laminar FSS.

In response to laminar FSS, we found that upregulation of TERT was required for endothelial elongation, flow alignment, and nuclear polarization. These observations identify TERT as another FSS-sensitive factor that dictates, in part, the endothelial phenotype in response to FSS. The classical function of TERT is restricted to the telomerase complex that prevents the cellular senescence typical of aging, particularly in highly replicating tissues ^48^. However, we found no change in endothelial telomere length over the time course of our study, suggesting the role of TERT in FSS is independent of its nuclear telomerase function. This contention is supported by studies identifying non-canonical functions of Tert in more quiescent tissues. For example, mice lacking Tert exhibit profound repression of PGC1α expression in the liver that mediates impaired mitochondrial biogenesis and function, including gluconeogenesis ^49^. Endothelial TERT undergoes nuclear export in response to oxidative stress ^50^, and localizes in mitochondria to protect mitochondrial DNA ^37,44^. This latter effect may be a consequence of its reverse transcriptase activity towards mitochondrial RNAs ^51^. More recently, TERT has been implicated in the switch between NO· -mediated and ROS-mediated vasodilation in the human microcirculation ^38,52^ characteristic of both aging and coronary disease. Whether the functions of TERT in our system are related to its nuclear or its mitochondrial localization is not yet clear, although the latter seems plausible since we found TERT was required for the endothelial mitochondrial responses to laminar FSS.

Our study identified TERT upregulation as key to the upregulation of HMOX1 with laminar FSS. This is a novel observation that adds to the longstanding knowledge that laminar FSS upregulates HMOX1 and other genes under the control of the antioxidant response element (ARE; ^53^ via activation of the Nrf2-Keap1 system ^54^. These data fit well with prior knowledge that one stabilizing component of laminar flow on the endothelium is the promotion of an antioxidant state and that HMOX1 upregulation is a key element in this process ^55^. Our data prompt new speculation that Tert could be an important component of the overall cellular ARE phase II detoxification responses. This speculation is consistent with prior observations of Tert nuclear export and activation in response to oxidative stress in the endothelium ^56^. The precise element(s) in the Nrf2-ARE pathway that are sensitive to TERT-mediated regulation is not yet clear, although cooperativity between KLF2 and Nrf2 has been described in the endothelium ^57^.

An important element of our study is the observation that HMOX1 upregulation is key to endothelial flow alignment and elongation in response to laminar FSS both *in vitro* and *in vivo*. This is a profound observation in that it adds a completely new consequence of HMOX1 in endothelial cell phenotype. The classic function attributed to HMOX1 is heme degradation, whereas, with laminar FSS, HMOX1 is thought to promote the antioxidant phenotype indicative of quiescent endothelium in atherosclerosis-resistant vascular sites ^55^ We observed co-localization of HMOX1 and TERT in the endothelium, suggesting some level of cooperativity. Both proteins are known to associate with the nucleus and mitochondria ^51,58^ and TERT association with the latter appears to enhance endothelial resistance to oxidative stress via reverse transcriptase activity ^59^. HMOX1 localization in the mitochondria is in association with biliverdin reductase and cytochrome P-450 reductase, suggesting its local function is in heme degradation ^58^. In contrast, HMOX1 nuclear localization has been associated with the upregulation of genes important for oxidative stress ^60^ Our data indicate that HMOX1 inhibition did not impact transcription of genes for PGC1α, KLF2, or TERT; however, some transcriptional role of HMOX1 in other laminar FSS-responsive genes cannot yet be ruled out. Nevertheless, our data provide a new role for HMOX1 in the endothelium, and the definition of the specific mechanism(s) involved will require further study.

## Methods

### Animals

C57BL/6J and VE-Cre (#006137) strains of mice were obtained from the Jackson Laboratories. PGC1α transgenic mice ^21^, conditional PGC1α ^61^, TERT global knockout ^62^ and conditional TERT ^63^ mice have been described previously. For the endothelial-specific knockout line, the PGC-1α or TERT allele containing *LoxP* sites flanking exons 3–5 of the PGC-1α gene (*PGC1a-floxed-* Bruce Spiegelman Harvard University) ^61^ or exons 1–2 of the TERT gene was bred with either VE-Cre or Tie2-Cre mouse line on the C57 background. This endothelial-specific PGC-1α (PGC1α ECKO) or TERT knockout mice were compared to Cre control mice. Endothelial specific PGC1a transgenic mice (PGC1α-ECTG) were produced with human PGC-1α expression under the control of the mouse vascular endothelial cadherin promoter (VE-Cad).

Exercise experiments were performed with 4-week old animals (PGC1α ECKO and Tie2Cre control mice), which were kept in individual cages for six weeks equipped with a running wheel and a mileage counter. Exercise training was performed voluntarily. The running difference in Tie2Cre control mice did not differ significantly from the PGC1α ECKO mice.

All mouse experiments have been done according to all the relevant ethical regulations. Mice were housed in a facility accredited by the American Association for Laboratory Animal Care and were granted by the Ethics Committee of the University Hospital Mainz and Landesuntersuchungsamt Koblenz (23 177-07/G 12-1-080 and 23 177-07/G 17-1-066). All animal studies were approved by the Institutional Animal Care and Use Committee of the Brigham Women’s Hospital, Boston.

### Cell Culture

Human Aortic ECs (HAECs) (#PCS-100-011) and Human Umbilical Vein ECs (HUVECs) (#PCS100010) cells were purchased from ATCC and cultured in EBM™-2 Endothelial Cell Growth Basal Medium-2 containing bullet kit growth factor supplements (Lonza), 5% fetal bovine serum, 100 units/mL Penicillin, 100 µg/mL Streptomycin, and 2 mM L-glutamine (Invitrogen). Cultured human ECs between passages 2 and 6 were utilized for experiments.

Adult mice sacrificed for MLECs isolation were anesthetized according to the approved guidelines mentioned above. Pooled mouse lungs were immediately harvested and stored on ice in a 50 mL conical tube containing Dulbecco’s Modified Eagle Medium (DMEM)(Gibco). Immersed mice lungs were then transferred to a 60 cm^2^ cell culture dish and chopped using sterile dissecting scissors. Minced tissues were transferred to a fresh 50 mL conical tube containing digestion solution, 50 mg/mL Type 1 filtered collagenase (Worthington) in DMEM media, and set to incubate at 37 °C for 1 hour. The minced tissue was filtered through a 70-μm and 40-μm cell strainer. The cell suspension was pelleted by centrifugation at 300g for 10 min at room temperature. The isolated cells were further used for sorting and direct mRNA isolation or for cell culture experiments.

For direct use of the cells in qPCR measurements, two separation steps were performed to isolate endothelial cells using CD31 MACS (CD31 Micro Beads mouse, 130-097-418, Miltenyi Biotech) beads and ICAM Dynabeads (Rat anti-mouse CD102/ICAM2, 553326, BD Bioscience) according to the manufacturer’s protocols.

For cell culture experiments, the pellet was resuspended in fresh MLEC growth media containing a 1:1 mixture of DMEM and Ham’s F-12 (Gibco), 20% Fetal Bovine Serum, 50 mg Endothelial Mitogen (Cell Applications), 50 mg/mL Heparin (Sigma), 100 units/ml Penicillin, 100 µg/ml streptomycin and plated on 0.2% gelatin-coated cell flask. Media was changed daily. When cell culture reached 80% confluency, cells were removed using Trypsin-EDTA (.05%) (Gibco). An equal volume of cell media was added, and cells were collected in a 15 mL conical tube with 25uL Dynabeads® sheep anti-Rat IgG (Invitrogen) coated with ICAM-2 (BD Biosciences Pharmingen) for the first selection. The cell suspension was set to rotate at 4 °C for

40 minutes. Once complete, the suspension was placed on a magnetic rack, and the supernatant was aspirated, and the bound beads were washed twice with MLEC growth media and once with DMEM media. The beads were resuspended in MLEC growth medium and plated into a new 0.2% gelatin-coated cell flask. When cells reached 80% confluency, steps were repeated for the second selection. Cells used for experiments were between passages 2-4.

### Adenoviral Constructs

An adenoviral vector expressing Adenovirus was used for both overexpression (24 hrs) and knockdown (48 h) of PGC1α (a kind gift from the laboratory of Bruce Spiegelman) and ERRα (a kind gift from the laboratory of Anastasia Kralli, Scripps Research Institute ^64^). Control viruses (siCtl, LacZ, and GFP) from Vector BioLabs (Malvern, PA). Cells were typically infected at a multiplicity of infection (MOI) of 10 to 50 with a control adenovirus at the same MOI.

### Transfections

Transfection assays were performed using 100nM small interfering human RNA oligonucleotides ON-TARGET plus SMART pool for control (D-001810-10), *TERT* (human-L-003547-00), *KLF4* (L-005089-00), *ERRa* (L-003403-00), and *HMOX1* (L-006372-00) (Horizon Discovery Dharmacon, Lafayette, CO) in DharmaFECT 3 reagent (Horizon Discovery Dharmacon, T-2003) for 6-8 hours in OptiMEMm (Invitrogen, Waltham, MA)^65^. Media was then changed to EBM™-2 Endothelial Cell Growth Basal Medium-2 containing bullet kit growth factor supplements (Lonza), 10% fetal bovine serum, 100 units/ml penicillin, 100 µg/ml streptomycin, and 2 mM L-glutamine (Invitrogen). For flow experiments, after 48 hours of siRNA treatment, cells were exposed to either laminar flow 12 dynes/cm^2^ or oscillatory flow of 4 dynes/cm^2^ (IBIDI system) for 48 hours.

### RNA Preparation and Quantitative Polymerase Chain Reaction

Cell and tissues were lysed with TRIzol reagent (Life Science Technologies), and total RNA was extracted using the RNeasy Plus Micro Kit (Qiagen). Total RNA was reverse transcribed with oligo(dT) primers for cDNA synthesis using an iScript cDNA synthesis kit (Bio-Rad). The expression of mRNA was examined by quantitative PCR analysis using a QuantStudio™ 6 Flex Real-Time PCR System (Applied Biosystems). TaqMan^©^ assays were used to quantitate *ERRα* (Hs00607062_gH), *HEY1* (Hs05047713_s1, Mm00468865_m1), *HMOX1* (Hs01110250_m1, Mm00516005_m1), *ICAM* (Hs00164932_m1), KLF2 (Hs00360439_g1, Mm00500486_g1), *KLF4* (Hs00358836_m1), *PGC1α* (Hs01016719_m1, Hs01016724_m1, Mm01208835_m1) *TERT* (Mm01352136_m1), *VCAM* (Hs01003372_m1), *UCP3* (Hs01106052_m1, Mm00494077_m1), *TBP* (Mm00446973_m1), *HPRT* (Hs02800695_m1, Mm00446968_m1), *GAPDH* (Hs99999905_m1, 4352339E-0904021) mRNA (Applied Biosystems), *B2M* (Hs99999907_m1, Mm00437762_m1). The 2^-ΔΔ*CT*^ method is used for relative quantification of gene ^66,67^. Reference genes of *HPRT, GAPDH, TBP and B2M* were used to normalize the PCRs in each sample.

### Immunoblot Analysis

Cell extracts were prepared using triton lysis buffer (TLB buffer) [20 mM Tris (pH 7.4), 1% Triton X-100, 10% Glycerol, 137 mM NaCl, 2 mM EDTA, 25 mM β-Glycerophosphate] with proteinase inhibitors (Sigma #11873580001) and phosphatase inhibitors (Sigma #4906837001). Protein extracts (50 µg of protein) in β-mercaptoethanol containing SDS sample buffer were separated in 4% to 12% gradient SDS-polyacrylamide gels (Bio-Rad #456-8094) and transferred to nitrocellulose membranes (Bio-Rad #170-4271, Hercules, CA) and incubated with a primary antibody with 1:1000 dilution. Immunocomplexes were visualized with horseradish peroxidase-conjugated secondary antibodies and detected with a Clarity Western ECL substrate (Bio-Rad #170-5061, Hercules, CA), and images were acquired on a chemiluminescent imager (Bio-Rad Chem-Doc Imaging System).

### Antibodies and reagents

Primary antibodies for immunoblots were obtained from Abcam (HMOX1 #52947, TERT # ab32020, Cambridge, MA), Cell Signaling (ERRα #13826, Danvers, MA); R&D (KLF4 #AF3640); Novus Bio (PGC1α #NBP-04676, Centennial, CO); BD Pharmingen (CD31 #550274, San Jose, CA) and Sigma (β-catenin, #C2206, St. Louis, MO). Antibodies used as controls were obtained from Proteintech (GAPDH #HRP-60004 and β-Actin #HRP-60008, Rosemont, IL). TERT inhibitor, BIBR1532 was purchased from Selleck Chemicals. HMOX1 inhibitor zinc (II) protoporphyrin IX (ZnPP) and control, copper protoporphyrin IX (CuPP)were purchased from Sigma. Another HMOX1 inhibitor 1-[[2-[2-(4-Bromophenyl)ethyl] -1,3-dioxolan-2-yl] methyl]-1*H*-imidazole hydrochloride (OB 24 hydrochloride) was purchased from Toris bioscience.

### Immunoprecipitation

Cell extracts were prepared using Triton lysis buffer [TLB - 20 mM Tris (pH 7.4), 1% Triton X-100, 10% glycerol, 137 mM NaCl, 2 mM EDTA, 25 mM β-Glycerophosphate, 1 mM sodium orthovanadate, 1 mM phenylmethylsulfonyl fluoride, and 10 µg/mL of aprotinin and leupeptin] and incubated (16 hrs., 4°C) with either 3 µg non-immune control rabbit IgG (Cell Signaling #272) or with 3 µg of ERRα antibody (Cell Signaling #13826) to 500 ul of cell lysate. Immunocomplexes were isolated using Protein G Sepharose beads (Santa Cruz SC2002, Dallas, TX) and 4-5 times washed with lysis buffer. Bead pellets were resuspended and boiled in β-mercaptoethanol containing Laemmli sample buffer were separated in 4% to 12% gradient SDS-polyacrylamide gels (Bio-Rad #456-8094) and transferred to nitrocellulose membranes (Bio-Rad #170-4271, Hercules, CA) and incubated with rabbit anti PGC1α antibody with 1:1000 dilution.

Immunocomplexes were visualized with horseradish peroxidase-conjugated goat anti-rabbit IgG (Cell Signaling, #7074, Danvers, MA) and detected with a Clarity Western ECL substrate (Bio-Rad #170-5061, Hercules, CA).

### Chromatin Immunoprecipitation (ChIP)

HUVEC cells were cultured and applied 1% oxygen for 30 min at 37C. ChIP kit was purchased from Active Motif (#53008, Carlsbad, CA), and the processing was followed according to the manual. Briefly, the cells were fixed in 1% formalin and homogenized in the lysis buffer. Lysed cells were sheared with sonication for ten times of pulse of 20 seconds with 30-second rest on ice between the shearing. Sheared chromatins were incubated with Protein G Beads and non-immune IgG (Cell Signaling #2729), Polymerase II (Santa Cruz #sc56767), PGC1a (Novus Bio #NBP-04676, Centennial, CO) or ERRa (Cell Signaling #13826, Danvers, MA) on an end-to-end rotator for overnight at 4C. The beads binding target chromatin were washed on a magnetic bar with washing buffer. Elucidated was target chromatin was amplified by nested PCR (Thermo #13001012) by using a primer pair 5’-CAGAAGTTTCTCGCCCCCTT-3’ and 5’-GAGGCCAACATCTGGTCAC-3’ then separated on 2% agarose gel. Approximately 200 kb bands were visualized with 0.5 ug/ml ethidium bromide and documented on a UV imager (Bio-Rad Gel Doc EZ imager).

### Whole Mount Aorta *En Face* Immunofluorescence Staining

Prior to excision, aortae were perfused via the left ventricle of the heart with 0.5 mM EDTA containing PBS solution and 4% Paraformaldehyde 7.5% sucrose 0.5 mM EDTA containing PBS solution and 0.5 mM EDTA containing PBS, respectively. Dissected aortae were fixed in 4% paraformaldehyde for 30 minutes and then washed with 0.1% Triton X-100 in PBS at room temperature, followed by incubation rabbit anti-β-Catenin antibody (Sigma #C2206, St. Louis, MO in blocking solution (Dako-Agilent, K800621, Carpinteria, CA) (dilution 1:1000) overnight at 4□°C. Repeating wash with 0.1% Triton X-100 in PBS, vessels set to incubate overate in secondary anti rabbit antibody conjugated with Alexa Plus 555 (Thermo #14-387-071, Waltham, MA)(dilution 1:1500) and nuclei were stained with DAPI for 5 minutes at room temperature. Vessels were mounted onto the coverglass with endothelial facing down and sealed on a coverslip with ProLong anti fade mounting medium (Thermo #P36962, Waltham, MA). Images were acquired with Confocal (Carl Zeiss) and ZEN 2012 software (Carl Zeiss).

### *In Vitro* Shear Stress to Endothelial Cells

*In vitro* endothelial cell fluid flow experiments were conducted using either of two different systems, the Ibidi system or a parallel plate flow chamber system previously described ^68^. The Ibidi system used endothelial cells from human and mouse seeded in μ-Slide I 0.4mm Luer, ibiTreat-tissue culture treated chambers (Ibidi #80176). Chambers for endothelial cells from mice were pretreated in 0.2% gelatin. After a confluent monolayer was reached, fluid flow conditions of either unidirectional steady flow (12 dyn/cm^2^), or bidirectional oscillatory flow (±4 dyn/cm^2^) were applied using the Ibidi Pump System (Ibidi #10902) for 48-hour treatment. The second system used a parallel plate flow chamber with a peristaltic pump that generates fully antegrade pulsatile flow (maximum, minimum and mean wall shear stress equal 6.7, 2.7, and 4.8 dyn/cm^2^, respectively) or net antegrade flow with a flow reversal component (maximum, minimum and mean wall shear stress equal 1.6, -1.1, and 0.3 dyn/cm^2^, respectively). The steady flow and the fully antegrade pulsatile flow waveforms are referred to as undisturbed flow (UF) since the flow is unidirectional. The bidirectional oscillatory flow and the net antegrade flow with a flow reversal component waveforms are referred to as disturbed flow (DF) because of the multidirectional nature of the waveforms. The fluidic units were maintained in 37 °C incubators with 5% CO2.

### MitoTracker Staining

Mitochondria in live cells were stained with either MitoTracker obtained from ThermoFisher (MitoTracker Red, M7512, Carlsbad, CA) or CytoPainter MitoRed from Abcam (#ab176832, Cambridge, MA) according to the manufacturer’s instructions. In Brief, the cells were incubated with 1:1000 diluted MitoTracker/MitoRed in a growth medium for 15 minutes at 37C incubator. Following the incubation, cells were washed in the growth medium twice and images were acquired.

### ROS Measurement

#### Amplex Red

As an index of ROS generation, we used the Amplex Ultra Red reagent, 10-acetyl-3,7-dihydroxyphenoxazine (Molecular Probes; A36006) that reacts with hydrogen peroxide (1:1 stoichiometry) in the presence of horseradish peroxidase (HRP) to form resorufin. Endothelial cells were cultured to confluency in 12-well plates and incubated with Kreb’s HEPES buffer (118mM NaCl, 22mM HEPES, 4.6mM KCl, 2.1 mM MgSO_4_, 0.15mM Na_2_HPO_4_, 0.41Mm KH_2_PO_4_, 5mM NaHCO_3_, 5.6mM Glucose, 1.5mM CaCl_2_) for 30 min. The Amplex Ultra Red and HRP were then added and fluorescence (excitation 544 nm; emission 590 nm) was determined as a function of time (2h) in 96-well black plates (Corning) at 37°C in a fluorescent plate reader (Spectramax, Molecular Devices). Vascular hydrogen peroxide formation in aortic tissue was measured by an HPLC-based Amplex Red assay as described earlier using aortic ring segments (3 mm in length) ^69^.

#### MitoSOX™ Red and DHE

Total cellular ROS and mitochondrial ROS levels were examined using DHE and MitoSOX™ Red, respectively, according to the manufacturer’s instructions. Briefly, cells were washed and loaded with DHE (10 µM) or MitoSOX™ Red (5 µM) at 37°C for 20 min in the dark and washed three times with MTB. Pre-loaded cells were incubated at RT, 37°C, 40°C or 42°C for 1, 2, or 3 h, respectively. To analyze ROS production in aortic tissue topographically and determine the source of ROS production, aortic cryo-sections were used as described previously ^70,71^.

### Immunofluorescent Staining

Cells were grown on #1 thick cover glass (EMS #72290-09, Hatfield, PA) or in flow chambers. Cells were fixed with 2% PFA or methanol for 10 min followed permeabilized with 0.1% Triton X-100 (Sigma Aldrich #X100) in PBS. Fixed cells were blocked in blocking solution (Dako-Agilent, K800621, Carpinteria, CA) and primary antibodies were incubated at 1:1000 overnight at 4 °C. Primary antibodies that were used are GOLPH4 #ab28049, HMOX1 #ab52947, TERT # ab32020, Cambridge, MA), Immunocomplex was visualized with anti-rabbit secondary antibody conjugated with Alexa Plus 555 (Thermo #14-387-071) or Alexa Plus 488 (A11034, Waltham, MA; dilution 1:1500). Autofluorescence was blocked and mounted with an antifade mounting medium (Vector Laboratories, Burlingame, CA).

### Quantification of Image

Using ImageJ processing software (National Institute of Mental Health, Bethesda), fluorescent images were imported, and individual color channels were separated. All area and intensity values were measured from the green channel. Under digital magnification, the width and height of each hyperfluorescent vessel were manually outlined and the encompassed measurement in pixels converted to μm using the scale bar. The average width/height ratio of experimental vessels was compared with the control vessels. For cell polarization, the locations of the nucleus and Golgi were identified, and the angle between the two points relative to the direction of flow was quantified. Three fields were used for quantification from each condition of the experiments. The wind rose plots were compiled using Origin software (OriginLab 2019, Northampton, MA).

### Isometric Measurements of Aortic Function

Thoracic aortic rings (2 mm in length) were mounted on 200 μM pins in a 6-mL vessel myograph (Danish Myo Technology) containing physiological salt solution (PSS): 130mM NaCl, 4.7 mM KCl, 1.18mM KHPO_4_, 1.17 mM MgSO_4_, 1.6 mM CaCl_2_, 14.9 mM NaHCO_3_, 5.5 mM dextrose, 0.03 mM CaNa_2_/EDTA. Vessels were stretched to 1g basal tension at 37°C and aerated with 95% O_2_-5% CO_2_. Vessels were equilibrated in PSS for 1h, followed by two consecutive contractions with PSS containing (60 mM potassium) and 1 μM phenylephrine (PE), then with KPSS alone. Rings were then washed, allowed to return to basal tension, and subjected to concentration-response curves to increasing concentrations of phenylephrine (PE), acetylcholine (Ach), and nitroglycerin (NTG).

### Telomerase Length

Telomere length was analyzed using a quantitative polymerase chain reaction (RT-qPCR)-based method previously described ^72,73^). The relative telomere length was calculated as the ratio of telomere repeats to a single-copy gene (SCG) (T/S ratio). The acidic ribosomal phosphoprotein PO (*36B4*) gene was used as the SCG. All qPCRs were performed in duplicates. The primers used for the telomere and the SCG amplification were as follows: telomere forward: 5′ GGT TTT TGA GGG TGA GGG TGA GGG TGA GGG TGA GGG T, telomere reverse: 5′ TCC CGA CTA TCC CTA TCC CTA TCC CTA TCC CTA TCC CTA; SCG forward: 5′ CAG CAA GTG GGA AGG TGT AAT CC and SCG reverse: 5′ CCC ATT CTA TCA TCA ACG GGT ACA A.

### Statistical Analysis

All data are expressed as mean ± SE, and the numbers of independent experiments are indicated. Statistical comparisons were conducted between 2 groups by use of Student t-test or Mann–Whitney U test as appropriate. Multiple groups were compared with either 1-way Kruskal–Wallis or ANOVA with a post hoc Tukey–Kramer multiple comparisons test as indicated in legends. A P value <0.05 was considered significant. All statistics were done using StatView version 5.0 (SAS Institute, Cary, NC) or GraphPad Prism version 5 (GraphPad Software, La Jolla, CA).

## Data Availability

The data sets generated and analyzed as part of this study are available upon request from the corresponding author.

## Author Contributions

S.K., K.V.T., M.K., S.K.S., J.M.J., S.M.C., C.S.T., E.S., and J.F.K. designed research. S.K., K.V.T., M.K., S.K.S., A.D.C., H.J.Y., S.M.C. and J.M.J. performed research. C.S. provided mice. S.K., K.V.T., S.K.S., J.M.J., S.M.C., C.S., E.S., and J.F.K. analyzed the data. S.K., A.D.C., K.V.T., S.K.S., J.M.J., S.M.C., H.J.Y., C.S., E.S., and J.F.K. wrote the paper.

## Acknowledgments

We thank Jennifer Cederberg, Jason Hagan, and Marisol Diaz for academic assistance; Claire C. Chu, Harish Janardhan, and Chinmay Trivedi for technical assistance; and Anastassiia Vertii for critical reading. We would like to thank the Microscopy Resources on the North Quad (MicRoN) Core staff for their training and support. This work was supported by grants 16SDG29660007 from AHA (to S.K.), CAREER CMMI1842308 from NSF (to J.M.J.) and K01AR073332 (to S.M.C.), HL142932 (to C.S.), 5T32HL120823 (to K.V.T.), HL151626 (to J.F.K.) from NIH.

