## Supplemental Figures for "PGC1α Regulates the Endothelial Response to Fluid Shear Stress *via* Telomerase Reverse Transcriptase Control of Heme Oxygenase-1"

**Supplementary Figures 1-4**

**Supplementary Figures**

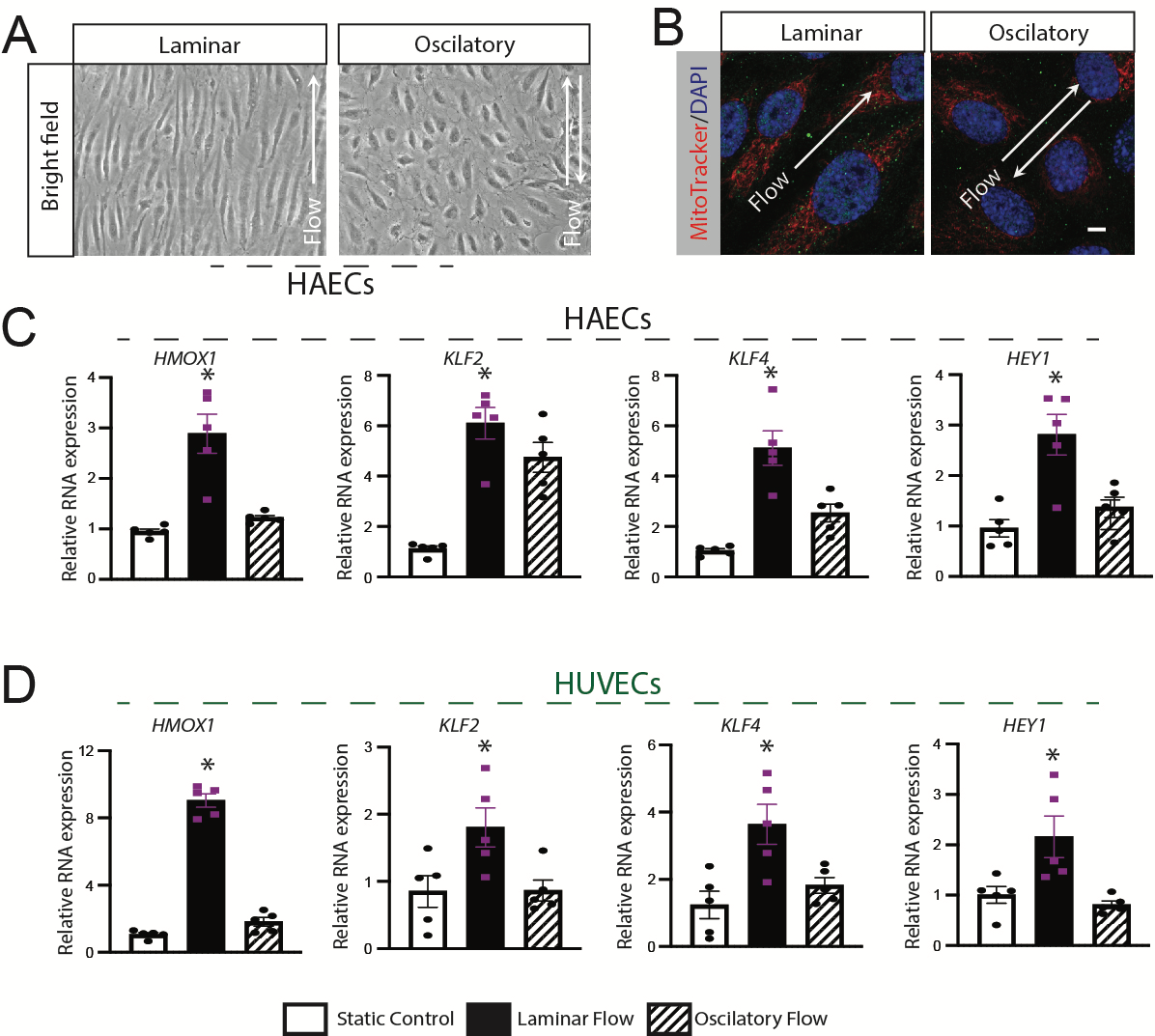

**Supplementary Figure 1. Endothelial cell alignment and gene expression during laminar shear stress.**

A) Alignment of human artery endothelial cells (HAECs) during flow shear stress (FSS) as described.

B) mitochondrial (red) and nuclear (blue) staining of HAECs during FSS. Scale bar, 5 μm.

C-D) Flow sensitive genes mRNA expression in HAECs (C) and human umbilical vein endothelial cells (HUVECs) (D) were measured by RT-qPCR after cells were harvested from either static, laminar, or oscillatory FSS for 48 hours (*n* = 4-5 in each group).**P* < 0.05 vs. indicated comparison by Student’s *t-test*. The data are mean ± SEM.

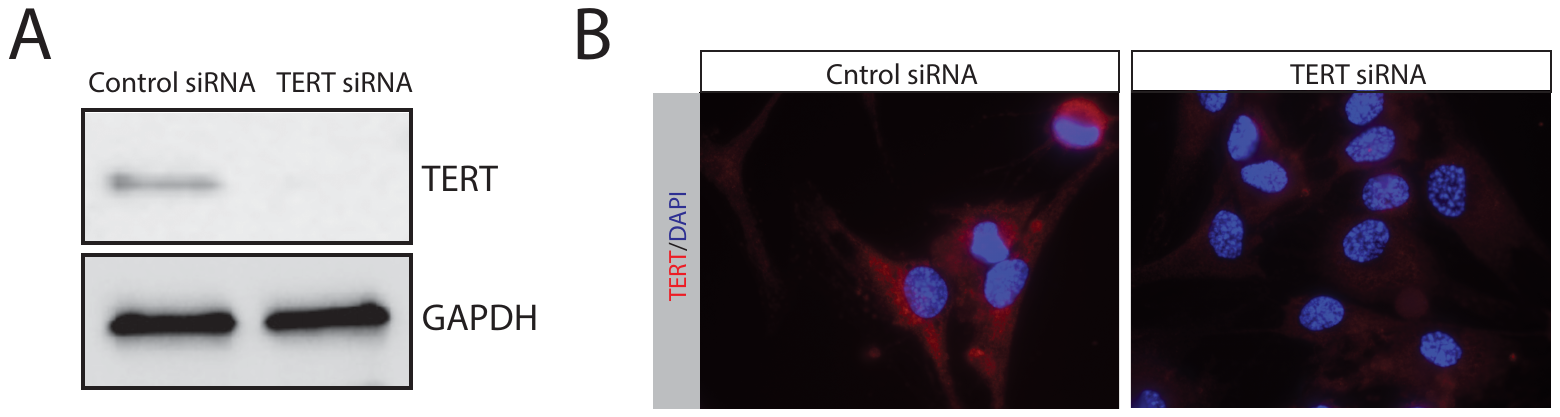

**Supplementary Figure 2.** A) Immunoblots for TERT and GAPDH antibodies were performed in HUVECs after the cells were treated for 48 hours with either control or TERT siRNA. B) HAECs were stained for TERT antibody (red) or Dapi (blue) after either treatment of control or TERT siRNA for 48 hours.

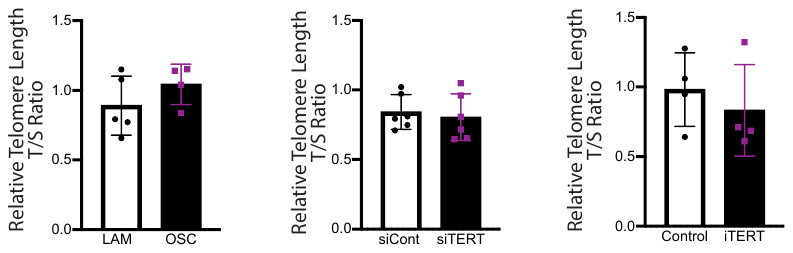

**Supplementary Figure 3.** Telomere length. HAECs telomere length was measured as a function of flow and either molecular (siTERT) or pharmacologic (iTERT) TERT inhibition. n=4-6.

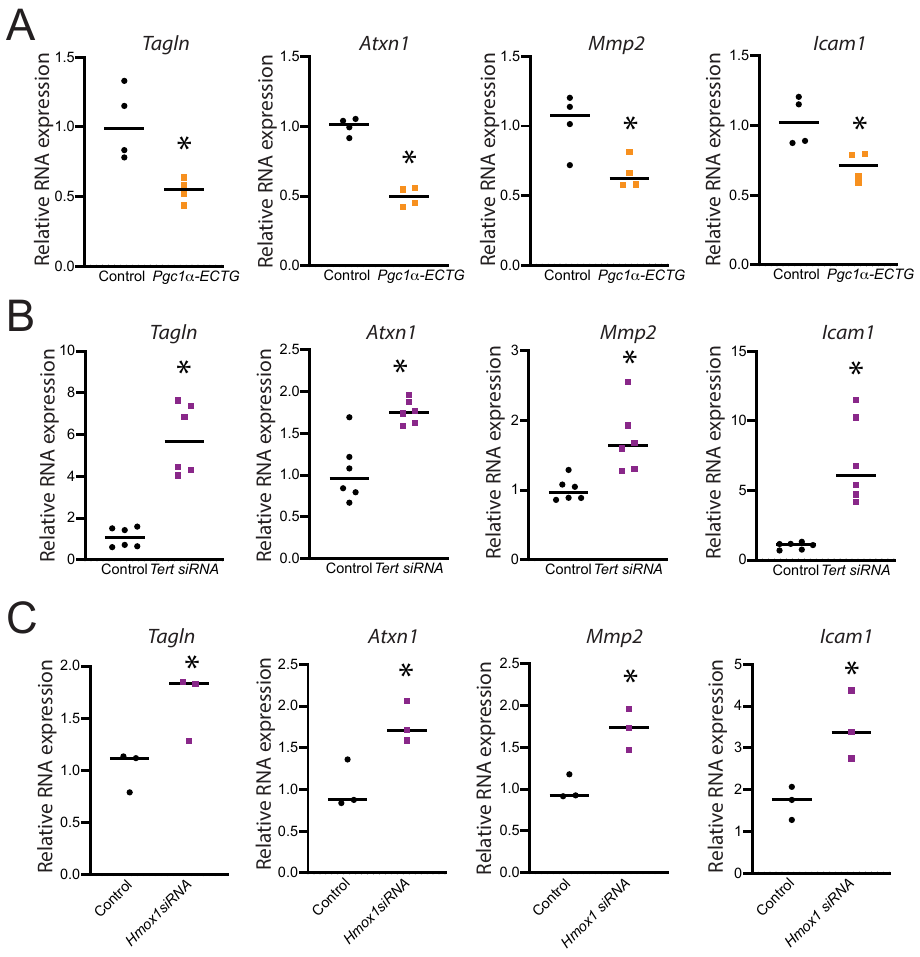

**Supplementary Figure 4. The PGC1α-TERT-HMOX pathway supresses endothelial to mesenchymal transition.**

A) MLECs were isolated from control and PGC1α -ECTG mice, and RT-qPCR was performed for mesenchymal markers SM22a (Tagln), Sca1 (Atxn1), Mmp2, and Icam1.

B-C) HAECs were either treated with scrambled or *TERT*siRNA (B) or scrambled or *HMOX1*siRNA (C), and RT-qPCR was performed for different mesenchymal markers (EndMT) after 48 hours. All the experiments have been repeated 3 – 6 times. Statistically significant differences between groups are indicated (**P* < 0.05 by Student’s *t-test*). The data are mean ± SEM. Scale bar, as indicated.
